# Bacterial communities associated with food-quality winter pea cultivars grown in Pacific Northwest soils

**DOI:** 10.1101/2024.11.25.625230

**Authors:** Svetlana N. Yurgel, Rebecca McGee

## Abstract

**Background and aims:** Breeding legume for improved yield and seed quality, coupled with extensive use of fertilization may disrupt the ability of modern cultivars to interact with native microbiome. Autumn-sown food-quality winter pea (WP) represent new crop in the Pacific Northwest (PNW). However, little is known about the ability of these cultivars to establish associations with bacteria native to PNW soils.

**Methods:** We evaluated soil, root and nodule microbiomes associated with four WP cultivars in diverse locations across Washington state with the goal to better understand the interaction occurring between WP cultivars and bacterial communities native to PNW soils.

**Results:** Root and nodule microbiomes were affected by sampling event, while plant genotype only affected nodule microbiome. A diverse population of native rhizobia colonized WP roots, while a smaller subset of these bacteria colonized WP nodules. Three rhizobial ASVs had relatively low abundance in the soils but were dominant in nodule-associated microbiome regardless of the variation of soil parameters between locations, indicating their strong attraction to host-plant nodules. Several non-rhizobial taxa were apparently enriched in nodules. However, in-depth study of legume root and nodule microbiome is required to better understand interactions within this complex phytobiome.

**Conclusion:** WP cultivars can form nodules in PNS soils in fall, but environmental factors have a strong effect on this process. While the complementation of legume nodule microbiome with root-associated microbiome analysis might be a useful tool, studies focusing on mature nodules with increased depth of sequencing might provide a better resolution of nodule-specific residents.

## 1. Introduction

There were 27,000 hectares of dry pea (*Pisum sativum* L) harvested in the US state of Washington (WA) in 2021 with a production value of $15 M (USDA, 2022). Peas are typically grown in rotations with small grain cereal, primarily wheat and barley. Although the great majority of peas are planted in the spring, there is increasing interest in using winter pea (WP) as a rotation, especially in the low-precipitation (<300 mm annual) region of the inland Pacific Northwest (PNW), where winter wheat is the dominant crop (Schillinger, 2017). WP is a dry field pea with cold tolerance suitable for autumn sowing in temperate regions and produces an early summer crop the following year. WP provides several advantages to spring-sown peas, including increased yield potential and seed-bed preparations in the fall, thus avoiding complications associated with cold, wet spring conditions. WP were primarily used for cover crop or feed purposes until 2009, when the Federal Grain Inspection Service permitted WP to be marketed as food-quality smooth dry yellow or green peas (USDA-GIPSA, 2010). Food-quality WP receive a premium price that is an economic benefit to growers. Several new food-quality WP cultivars are already in production. While analysis of WP food quality and management is ongoing, little is known of the ability for WP cultivars to establish symbiotic associations with native rhizobia in PNW soils.

Legume domestication and intensive selection for improved yield and seed quality, coupled with extensive use of nitrogen (N) fertilization may disrupt nodulation rate, N fixation, and/or host N uptake, as well as the ability of the legume to select for the most efficient rhizobial partner (Porter & Sachs, 2020). It was proposed that wild legume species develop molecular mechanisms to preferentially select for more mutualistic symbionts with higher N fixation potential (Younginger & Friesen, 2019). It was also predicted that legume domestication and breeding for high input farming systems might have inadvertently produced legume genotypes which are more dependent on soil N than on symbiotic nitrogen fixation. This can lead to alteration of host-bacteria compatibility, plant-associated microbiomes and the ability of the host to select for highly efficient bacterial partners (Turcotte *et al*., 2017, Liu *et al*., 2020, Porter & Sachs, 2020, Rahman *et al*., 2023). For example, it was reported that less than one half of native *Rhizobium leguminosarum* isolates capable of establishing effective symbioses with wild legumes were also able to establish effective symbioses with the domesticated species, demonstrating modern cultivars might be losing their interaction with rhizobia as compared with wild legumes (Mutch & Young, 2004). Similarly, it was shown that chickpea and wild soybean can recruit more diverse rhizobial partners in the nodules and rhizosphere than the cultivated species (Kim *et al*., 2014, Chang *et al*., 2019).

Symbiotic nitrogen fixation takes place in the organs formed on the host-plant roots. Most of pea nodulating rhizobia belong to the *Rhizobium leguminosarum* complex, defined as a group of closely related species, including *R. ruizarguesonis* and *R. laguerreae* (Young *et al*., 2021), as well as more the distant species, such as *R. pisi* (Ramírez-Bahena *et al*., 2008), *R. indicum* (Rahi *et al*., 2020) and *R. anhuiense* (Zhang *et al*., 2015). Additionally, peas form indeterminate nodules, where persistence of the meristem leads to the development of four functional zones: distal zone I with nodule meristem, zone II with infection threads occupied by bacteria, zone III with mature nitrogen-fixing rhizobia, and senescence zone IV (Poole *et al*., 2018). Rhizobia are not the only inhabitants in legume nodules (Martínez-Hidalgo & Hirsch, 2022). Other nodule-occupying bacteria originate from the soil and interact with the nodule microbiome to affect N fixation and fitness of the host plant. Therefore, the senescence zone IV of mature nodules might be colonized by non-rhizobial bacteria. The non-rhizobial bacteria are outnumbered by rhizobia but exhibit a striking level of diversity and have potential to benefit the host plant, especially under environmental stress. However, prolonged existence of nodules in such diverse ecological niche as soil makes it difficult to identify true non-rhizobial nodule endophytes by Next Generation Sequencing. In this study we profiled the root-associated microbiome to identify non-rhizobial bacteria enriched in WP nodule-associated microbiome.

This task was coupled with the broader goal of understanding the interaction occurring between WP cultivars and bacterial communities native to PNW soils. This was especially important since WP is planted in fall, and therefore it has a potential to form nodules before the low winter temperature prevents the nodulation. In this study we evaluated bacterial root and nodule microbiomes associated with three new food-quality WP cultivars and with one more genetically distant WP genotype grown in three diverse locations across WA. To assess the progression of WP bacterial community over winter, the microbiomes were collected in fall, two months after WP planting, and in early summer before WP flowering. To our knowledge this is the first spatiotemporal analysis of WP root- and nodule-associated microbiomes.

## 2. Materials and Methods

### 2.1. Site description

For this study, field trials at three locations (Pullman, Garfield, Dayton) in Washington were established in October 2021 (Table S1). All locations were planted to wheat and were sprayed with Dual Magnum selective herbicide at 1471 g/h (Syngenta, US), Roundup RT3 (Bayer Crop Science, US) at 3362 g/h, and Sharpen (ASF the Chemical Company) at 140 g/h for pre-emergence control in the fall of 2021. All locations received 701 g/h of selective herbicide Assure II (Keystone Pest Solutions, LLC) and 911 g/h of Crop-oil (Monterey) for grass control in the spring of 2022. All locations were sprayed with Warrior II (Zeon Technology) at 140 g/h and Downrigger (The McGregor Company, US) at 630 g/h for pea seed weevil. None of the location received any fertilizers or rhizobium inoculums. The data for weather condition for the sites are represented in Table S1 and Fig. S1. At each location four WP cultivars were planted, which include three new food-quality WP cultivars, ‘USDA MiCa’ (PI 699929), ‘USDA Klondike’ (PI 699931) and ‘USDA Dint’ (PI 699930) and a feed-quality WP cultivar ‘Windham’ (PI 347868). Windham was included in the study as a more genetically distant WP genotype in order to better evaluate the effect of seed-quality directed breeding on the plant-associated microbiome. Prior to planting, the pea seeds were treated with a slurry that included the fungicides Fludioxoinil (0.93 g kg−1; Albaugh, LLC, Ankeny, Iowa, United States), Rizolex (0.93 g kg−1; Albaugh), Mefenoxam (0.93 g kg−1;Albaugh), and Intego Solo (0.18 g kg−1; Valent, United States), the insecticide thiamethoxam (0.80 g kg−1; Albaugh), and Red Colorant (0.51 g kg−1). The slurry was applied at 4 ml kg-1 rate. The experiment was conducted in small plots (6 m × 1.5 m) planted in a randomized complete block design with three replications.

### 2.2. Nodulation test

Seeds were initially washed with water containing a small amount of Alconox detergent for 1 min, followed by 95% ethanol for 1 min and 15% hydrogen peroxide for 1 min. The seeds were rinsed 5 times with ultrapure sterile water between each treatment and after final treatment. For germination, both sterilized and non-sterilized seeds were soaked in ultrapure sterile water for 24 hours, then placed on agar plates and germinated in the dark at room temperature for 24-48 hours (until the radicle was between 1 and 2 cm long). Scott trifold towels were folded into thirds and tightly rolled before placing in the bottom of a 250 mm long glass culture tube (cat. 14-923R, Fisher Scientific), with caps and autoclaved. 25mL plant nutrient solution (PNS, Table S2) was added to each tube to completely wet the paper towel. A small pocket was made by pulling the paper towel away from the wall of the tube, and one sprouted pea planted in each tube by tucking it between the paper towel and the edge of the tube with the radicle pointing down. Aluminum foil was wrapped around the tube to 1cm above the top of the paper towel. The tubes were left on a benchtop for 2-3 days before inoculation. A moderately thick suspension (OD_600_ = 1) of *R. leguminosarum* Rl3841 (Johnston & Beringer, 1975) was made in sterile PNS, and 1mL of suspension was added to each tube, aiming for the pea root and tilting the tube if needed to make sure the bacteria come in contact with the root. 1mL PNS was added to uninoculated plants. The tubes were placed in a growth chamber (set to 22°C and 16 hours light/8 hours dark) and the nodulation was assessed after 5 weeks.

### 2.3. Plant tissue collection

The plant and soil samples were collected in December 2021 (fall collection) and late May 2022 (summer collection) Table S1. Plants, with attached root system, were carefully removed from the soil using shovel, placed in sterile bags, and transported to the laboratory on ice. At each sampling event, four to five plants from each plot were collected resulting in 16-20 plants for each cultivar. The roots with attached nodules were washed three times with 10% glycerol and sonicated three times as described previously [37]. Since all sterilization methods of nodule tissue are aimed to remove bacteria from nodule surface and none of the methods were proven to remove DNA from the surface [38], we used sonication step to remove most of rhizoplane community intact [39] leaving bacteria that are strongly attached to the root and nodule surface. This approach gave a better resolution of WP microbiome intimately associated with the plant root system. Only nodulated plants were used in further analysis. The nodules and roots were separated, washed in 10% glycerol, placed into cryogen vials, and stored at -80°C prior to DNA isolation. The frozen roots were ground into fine powder in liquid nitrogen and 0.25 g of root tissue was set aside for DNA extraction. Nodules were crushed using Disposable Pellet Pestles (Fisherbrand) and 0.1g of the tissue was used for DNA extraction.

### 2.4. Soil collection

During summer soil sampling, one soil sample was taken from each plot (12 samples per location, 36 samples total). Soil samples were immediately placed on ice and transported to the laboratory. The soils were sifted through a 2 mm sieve before being transferred into a -80°C freezer. For each sample, 0.250 g (wet weight) of sifted soil was used for DNA isolation.

### 2.5. Soil chemical analysis

Soil chemical parameters from fall collected soils were determined using Best Test Analytical Services, LLC (3394 Bell Rd NE Moses Lake, WA 98837). Each soil sample was characterized by determining bulk density (g/ml), pH, electrical conductivity (dS/m), organic matter (OM, %), total nitrogen (N, %), and total carbon (C, %) according to standard procedures.

### 2.6. DNA extraction and sequencing

Plant and soil and DNA extraction was carried out using the DNeasy PowerSoil Pro Kits extraction kit according to the manufacturer’s protocol (Qiagen). DNA quality and concentration were measured using NanoDrop® ND-1000 UV-Vis Spectrophotometer (Thermo Scientific). At least 50 ng (10 µL) of DNA samples were sent to the Dalhousie University Centre for Comparative Genomics and Evolutionary Bioinformatics–Integrated Microbiome Resource (CGEB-IMR) for V6–V8 16S rRNA gene (16S; forward: 5′-ACGCGHNRAACCTTACC-3′; reverse: 5′-ACGGGCRGTGWGTRCAA-3′). Samples were multiplexed using a dual-indexing approach and sequenced using an Illumina MiSeq with paired-end 300 + 300 bp reads. All PCR procedures, primers, and Illumina sequencing details were as described (Comeau *et al*., 2017).

### 2.7. Amplicon Sequence Processing

The sequence processing was performed using the protocol outlined in the Microbiome Helper package (Comeau *et al*., 2017). The reads were processed with QIIME2 (version 2020.8) (Bolyen *et al*., 2019). Sequences were filtered for low-quality or probable chimeric reads from the dataset by using QIIME2’s q-score-joined function. Using QIIME2’s Deblur plug-in, the sequences were organized into amplicon sequence variants (ASVs) high-resolution genomic groupings (Amir *et al*., 2017). Taxonomic classifications were assigned to the ASV using QIIME2’s naive-Bayes approach implemented in the scikit learn function, referencing SILVA databases (Bokulich *et al*., 2018). The low-abundant ASVs were removed, and ASVs assigned to mitochondria and chloroplasts were filtered out.

### 2.8. Data Analysis and Statistics

#### Plant-associated dataset

1,115,828 high-quality non-chimeric reads were obtained from plant 16S rRNA amplicon sequencing. These sequences were clustered into 2,239 ASVs. Four samples with total read < 2,169 were removed from the downstream analysis. For diversity assessment, the dataset was normalized to the depth of 2,169 reads. *Soil dataset:* 342,882 high-quality non-chimeric reads were obtained from soil 16S rRNA amplicon sequencing. These sequences were clustered into 9,979 ASVs. For diversity assessment, the dataset was normalized to the depth of 2,152 reads. Alpha-diversity (Chao1 richness, Simpson evenness and Shannon diversity) and beta-diversity metrics were generated using QIIME2 diversity core-metrics-phylogenetic plugin. The differences in microbial alpha-diversity were calculated using Kruskal-Wallis pairwise test (p < 0.05) with Ben-jamini-Hochberg FDR multiple test correction. Variations in sample groupings explained by Bray-Curtis beta-diversity distances (PERMANOVA/Adonis tests, 999 permutations) were run in QIIME2 to calculate how sample groupings were related to microbial community structure. Non-metric Multidimensional Scaling (NMDS) plots were built based on Bray-Curtis distances using the Vegan package in R (Oksanen *et al*.). NMDS plots were visualized using the gglpot2 package in R (Wickham, 2016). Differential relative abundances were determined using Analysis of Compositions of Microbiome (ANCOM) R package with 1% FDR (Mandal *et al*., 2015) and ANOVA-Like Differential Expression (ALDEx2) (Fernandes *et al*., 2013).

## 3. Results

### 3.1. Site weather conditions and soil chemical properties

The three locations differed in weather conditions and soil chemical composition. Dayton soil was significantly lower in organic matter and total nitrogen and carbon, compared to that in Garfield and Pullman. The Garfield soil had the highest electrical conductivity and organic matter and total carbon content (Table 1). All soils had relatively low pH with the highest pH of 5.37 found in Dayton. On average monthly air and soil temperature was the highest in Dayton area (Table S1, Fig. S1). Dayton also received the most precipitation in spring 2022 and Pullman received the most precipitation in fall 2021.

**Table 1.** Soil chemical properties. For each variable, data followed by different letters are significantly different according to Tukey’s test (P < 0.05)

| Parameters | Dayton | Garfield | Pullman |
| --- | --- | --- | --- |
| Bulk density, g/ml | 1.24<sup>A</sup> | 1.25<sup>A</sup> | 1.26<sup>A</sup> |
| Electrical conductivity, dS/m | 0.242<sup>B</sup> | 0.383<sup>A</sup> | 0.276<sup>B</sup> |
| Organic matter, % | 2.40<sup>C</sup> | 3.49<sup>A</sup> | 2.67<sup>B</sup> |
| pH | 5.37<sup>A</sup> | 4.98<sup>B</sup> | 4.95<sup>B</sup> |
| Total nitrogen, % | 0.098<sup>B</sup> | 0.123<sup>A</sup> | 0.128<sup>A</sup> |
| Total carbon, % | 0.97<sup>C</sup> | 1.44<sup>A</sup> | 1.18<sup>B</sup> |

### 3.2. Assessment of WP nodulation

A question of inconsistency of plant-microbe interaction and legume symbiotic performance between green-house and field condition remains not well answered. However, recent study comparing phytobiomes formed with *Lupinus angustifolius* plant grown in green-house and field conditions indicated that the bacterial nodule microbiome was highly influenced by the culturing conditions (Ortuzar *et al*., 2024). The study suggested that in the greenhouse, the plant filtering is less efficient than in the field. Furthermore, WP production cycle includes an extended period (3-4 month) of plant growth under low soil temperature and it is very difficult to reproduce these environmental conditions in greenhouse, making field experiments a better approach to study WP associated microbiome. However, it was not feasible to surface sterilize WP seeds used in the field experiments. Additionally, the interiors of the legume seeds might also contain rhizobial strains that can be propagated vertically and their ability to from nodules in the presence of exogenous rhizobia could not always be controlled by seeds sterilization (Mora *et al*., 2014). Therefore, we evaluated the ability of rhizobia carried by the seeds used in field experiments to form nodules. Non-sterilized seeds of four WP cultivars were propagated in sterile glass tubes for six weeks and nodule development was monitored. To ensure that the test conditions were suitable for WP nodulation, inoculated plants were used as a control. Many plants produced from non-sterilized seeds had fungal contamination and were removed from the analysis. At least 15 non-contaminated plants in each test group were scored for nodulation. None of the non-inoculated plants formed nodules. On the other hand, at least 47% of inoculated plants formed nodules (Fig. 1). This data indicated that the majority of the nodules formed on the WP roots grown in the field conditions were induced by native soil rhizobia.

**Fig. 1.**
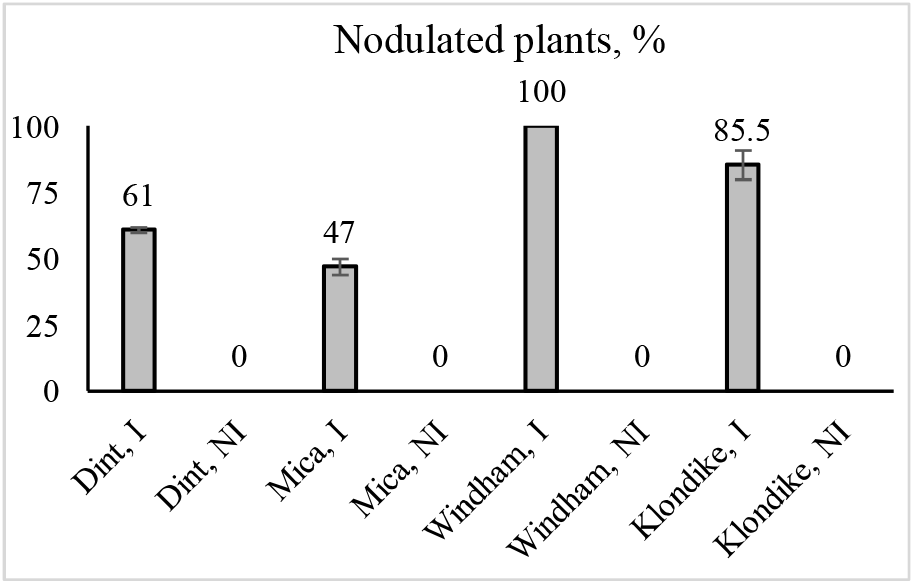
Ability of rhizobia carried by the seeds to form nodules. I – inoculation; NI – no inoculation. At least 15 seeds were tested in each treatment group in two biological replicas.

The timing for fall sample collection was aimed to provide the plants with time to establish nodulation, but prior to soil freezing, which could complicate sample collection. The summer collection time targeted the plant pre-flowering stage to avoid shutdown of N-fixation and associated rapid nodule senescence. However, at one location (Dayton), at the time of summer nodule collection, all nodules were destroyed by pea leaf weevil. Therefore, only nodules developed by December were collected at Dayton.

The plants grown in Pullman did not develop early nodules by December 2021 and only summer-collected plant tissue was used in the study. Approximately 50% of the plants collected in both summer and fall contained pink nodules (Table S1) and in most cases the nodules were relatively small. Therefore, the nodules from individual plants from the same plot were combined to compose a single sample with a mass of at least 0.5g. The list of samples is presented in Table S3. In total, 72 nodule and corresponding root samples were collected and sent for sequencing.

### 3.3. Overall WP microbiome

In total, 2,238 16S rRNA amplicon sequence variants (ASVs) were identified in this study (Table S4). Among them 549 ASVs, representing around 25% of all detected ASVs, were identified in both the root- and nodule-associated microbiomes, 1678 ASVs (75%) were identified only in roots and 11 were only found in the nodules (Fig. 2 A). We identified 120 unique ASVs annotated as *Allorhizobium-Neorhizobium-Pararhizobium-Rhizobium (Rhizobium*) with 65 of those were found both in roots and nodules. Three (781e9f4836b615ff81ab450ca826ec80, 840ed78ed13d6351d9f35625b150b98f, and bcdc13616a908c20ccc12b426b58feb6) were unique to the nodules (Fig. 2 B). *Rhizobium* was the most abundant genus identified in both root- and nodule-associated microbiomes and was represented by 78% and 99.5% of total root and nodule reads, respectively.

**Fig. 2.**
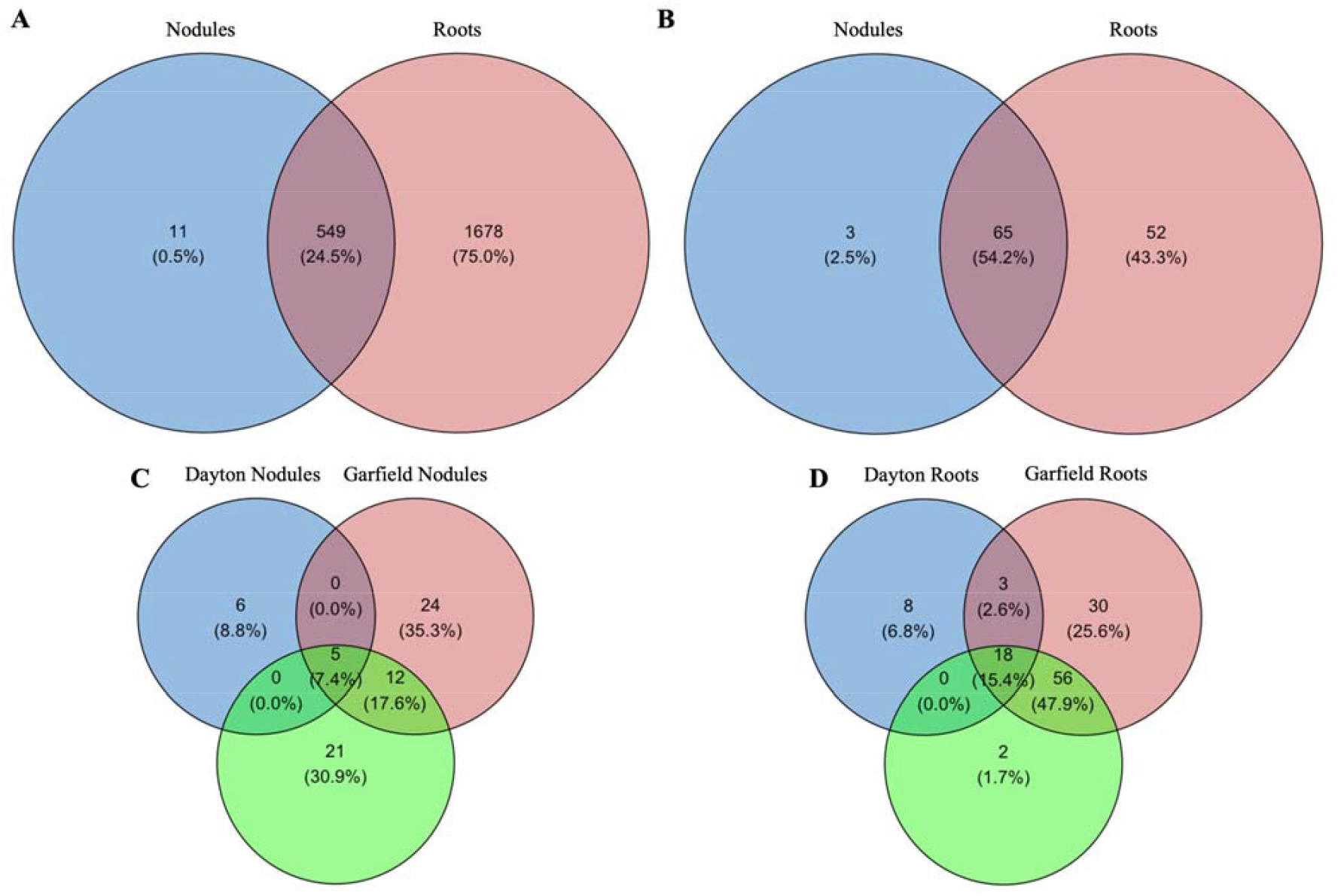
Venn diagrams showing overlap in ASVs between niches. Overlap in A – total and B – rhizobial ASVs found across the WP nodules and roots; and overlap in C – nodule and D – root rhizobial ASVs found between across locations

Cultivar was a significant but minor factor shaping total WP microbiome, explaining 2.5% of total WP community variation. When nodule- and root-associated microbiomes were considered individually, a significant effect of cultivar (9%) was only detected on the nodule microbiome (Table 1). When microbiomes were grouped together by cultivar or by cultivar and tissue type, no significant differences in their alpha-diversity were detected, which was consistent with the minor effect of this factor on bacterial beta-diversity. Additionally, we did not identify any bacterial taxa in total and tissue specific microbiomes differentially represented between WP cultivars.

Tissue type (root vs. nodule) was a strong factor affecting total WP microbiome, contributing 11.9% of the community variation (Table 2; Fig. 3). Furthermore, the nodule microbiome exhibited significantly lower Shannon diversity compared to the root, 1.6 vs. 3.7 respectively (Fig. 4). *Rhizobium* was the only genus significantly overrepresented in nodule, compared to root microbiome, while many genera outside the *Rhizobium* group were overrepresented in the roots (Table S5). More specifically, three ASVs c9d76e2a4779608de747a03a10b10649 (ASV1), 503453882a172c31ac560442c1d151cc (ASV2) and 9600d5ba182b6d29415b446a045ed0d1 (ASV3), which were manually annotated as *Rhizobium leguminosarum*, were significantly overrepresented in the nodules compared to the roots (Fig. 5 A, B, C). However, these ASVs were also abundant in root tissue and were represented by 95% and 63% of total nodule and root reads, respectively. These ASVs represented the core root and nodule microbiome – these were the only ASVs identified in 100% of nodule and 90% of root samples.

**Table 2.**
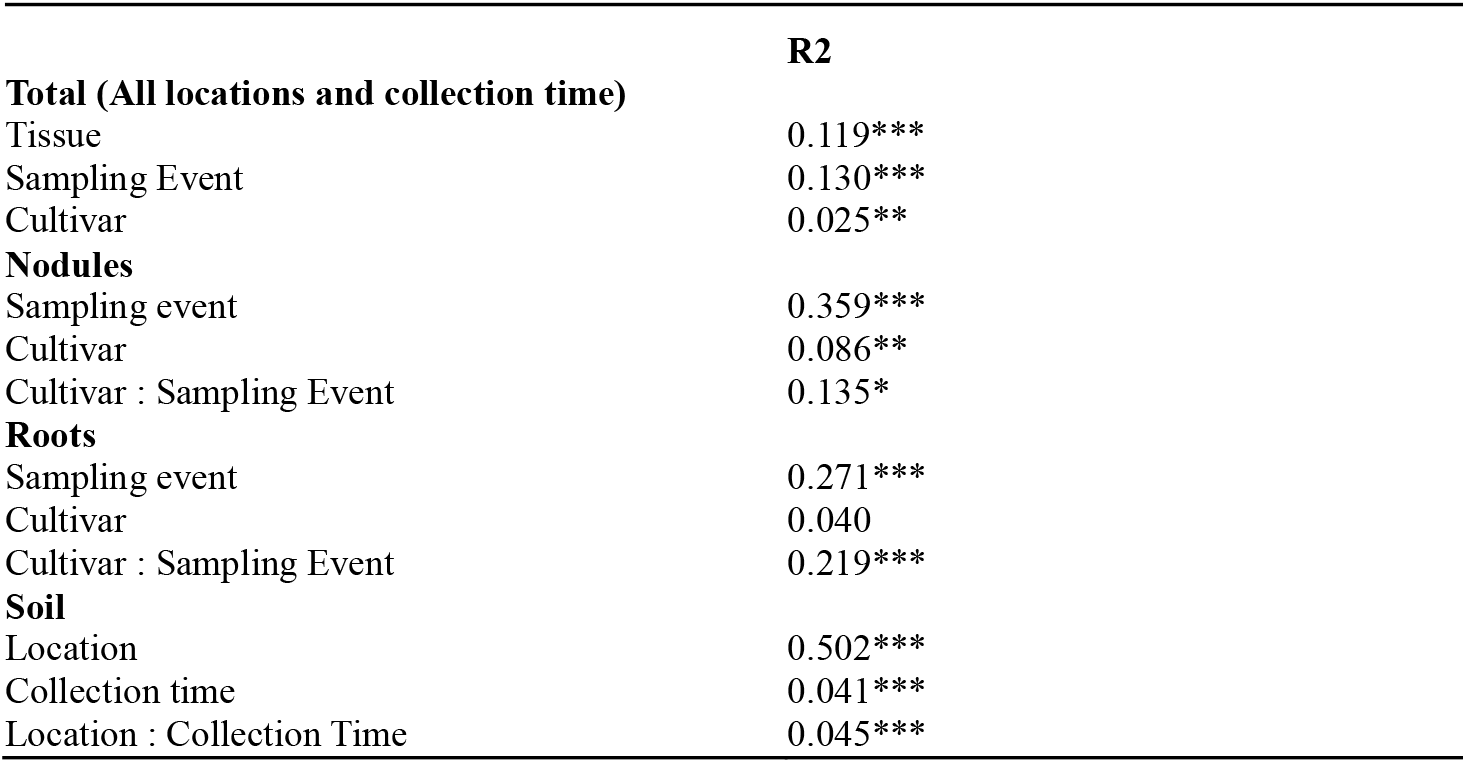
Variation in sample groupings as explained by weighted Bray-Curtis dissimilarity distances. Adonis tests were used to assess whether beta-diversity is related to sample groupings, 999 permutations, R2, *P < 0.05, **P< 0.01, and ***P < 0.001

|  | R2 |
| --- | --- |
| **Total (All locations and collection time)** |  |
| Tissue | 0.119*** |
| Sampling Event | 0.130*** |
| Cultivar | 0.025** |
| **Nodules** |  |
| Sampling event | 0.359*** |
| Cultivar | 0.086** |
| Cultivar : Sampling Event | 0.135* |
| **Roots** |  |
| Sampling event | 0.271*** |
| Cultivar | 0.040 |
| Cultivar : Sampling Event | 0.219*** |
| **Soil** |  |
| Location | 0.502*** |
| Collection time | 0.041*** |
| Location : Collection Time | 0.045*** |

**Fig. 3.**
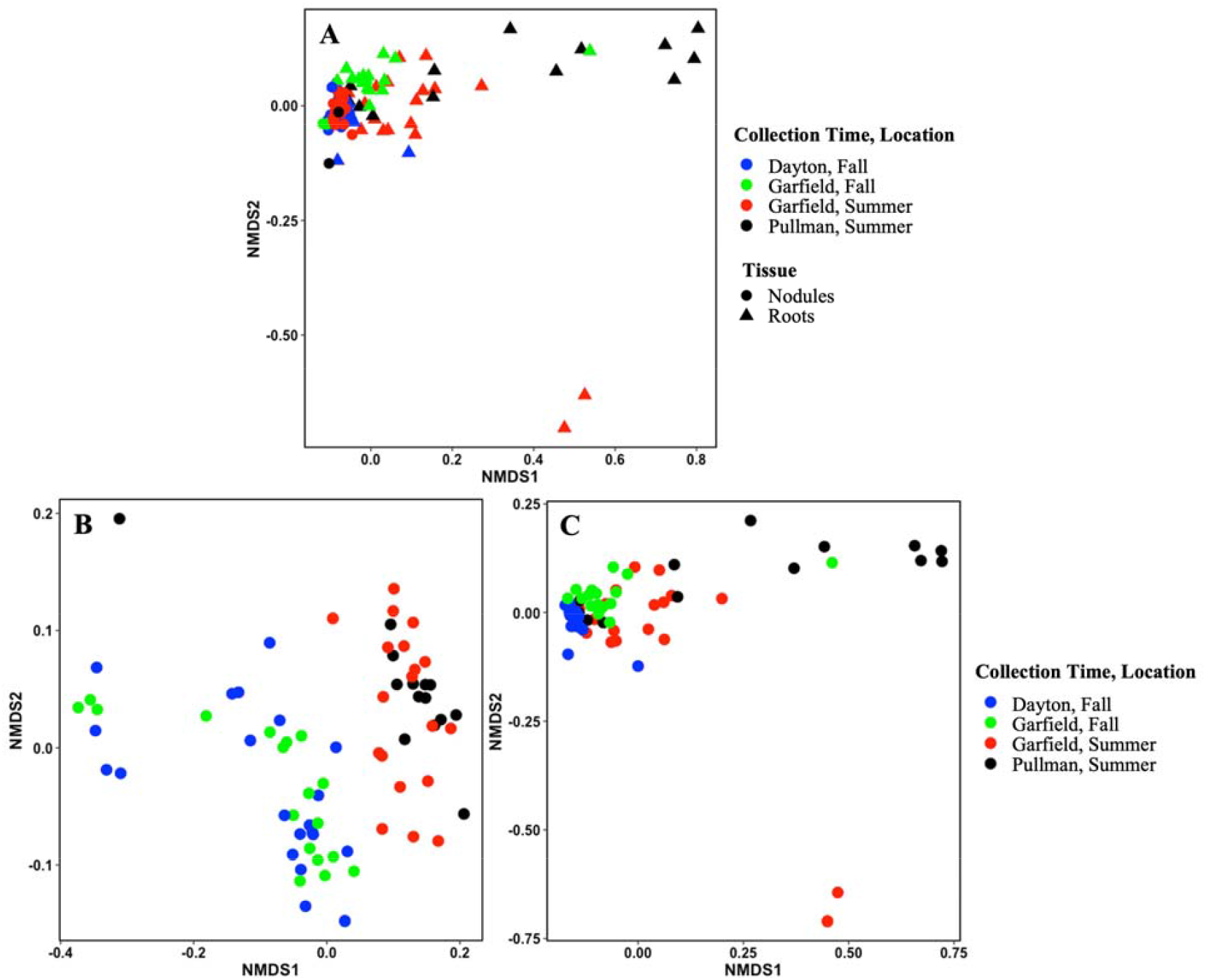
Bacterial beta-diversity in plant associated microbiome. Nonmetric multidimensional scaling (NMDS, based on Bray-Curtis dissimilarity distances) of bacterial communities at ASV level. A– root and nodule; B – nodule; and C - root microbiome

**Fig. 4.**
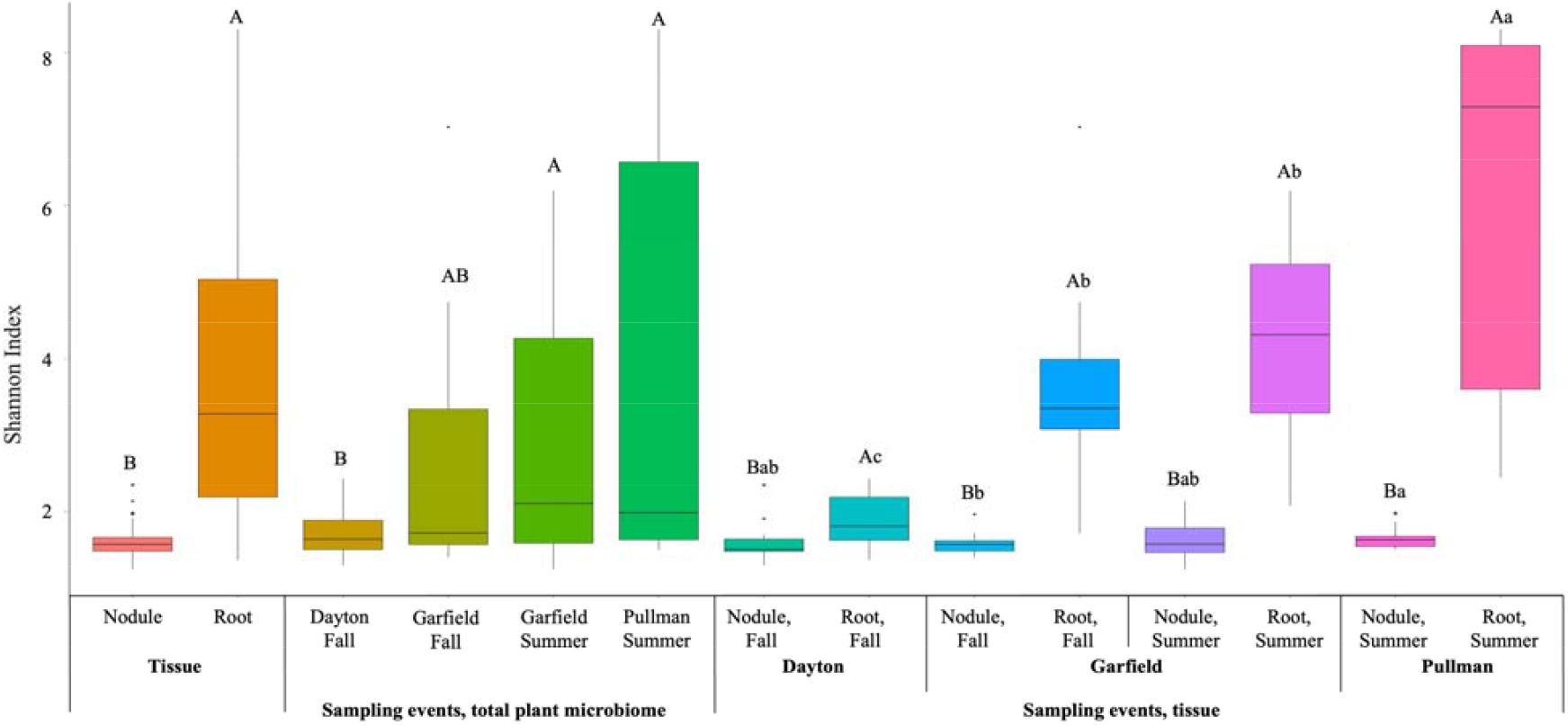
Estimated Shannon diversity of bacterial community. For each variable, data followed by different letters are significantly different according to Kruskal-Wallis pairwise test (p < 0.05). Corrected p-values were calculated based on Benjamini-Hochberg FDR multiple test correction. Capital letters indicate differences within each group. Lowercase letters indicate differences between sampling events for each tissue type

**Fig. 5.**
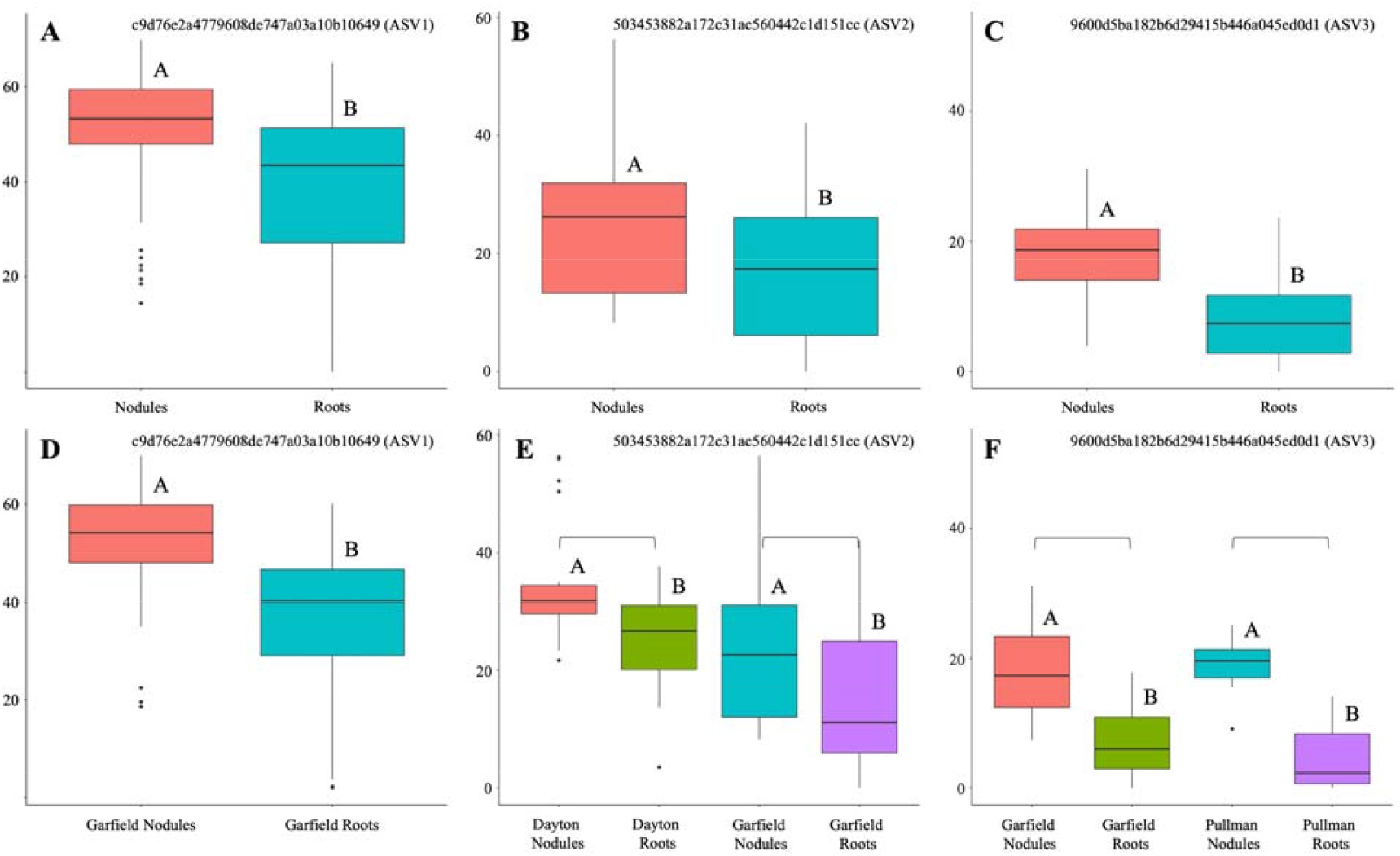
Relative abundances of ASVs differentially represented between nodule and root tissue. For each variable, data followed by different letters are significantly different according to Kruskal-Wallis pairwise test (p < 0.05). A, D – ASV1; B, E – ASV2; C, E – ASV3. A, B, C – relative abundances of ASVs in total nodule and root microbiome; D, E, F - relative abundances of ASVs across locations. Corrected p-values were calculated based on Benjamini-Hochberg FDR multiple test correction. Capital letters indicate differences within each group

#### 2.3.1. Non-rhizobial WP microbiome

To better resolve the non-rhizobial WP microbiome, ASVs annotated as *Rhizobium* were filtered out and the remaining microbiome was evaluated. However, after filtering rhizobial ASVs, 24 samples did not contain any reads and 48 samples had < 100 reads, resulting in a dataset with nine nodule samples and 63 root samples with at least 100 reads (Table S3). The *Gammaproteobacteria, Pseudomonas*, was the most abundant genus in the nodule-associated microbiome and was represented by approximately 27% of the total non-rhizobial nodule reads (Fig. 6 A; Table S6). While it was not statically significant *Pseudomonas* was 10 times more enriched in the nodules, compared to roots. Additionally, two Actinobacteria, *Leifsonia* and *Microbacterium*, were enriched in the nodules with their relative abundance 12 and 15 times higher in nodule than in root tissue, respectively. The other relatively high abundant bacteria in nodule-associated microbiome included *Tardiphaga, Mesorhizobium, Mycobacterium, Mucilaginibacter*, and *Flavobacterium*, which were represented by at least 4% of total non-rhizobial nodule reads. However, these genera were less relatively abundant in the nodule-associated microbiome compared to *Pseudomonas*.

**Fig. 6.**
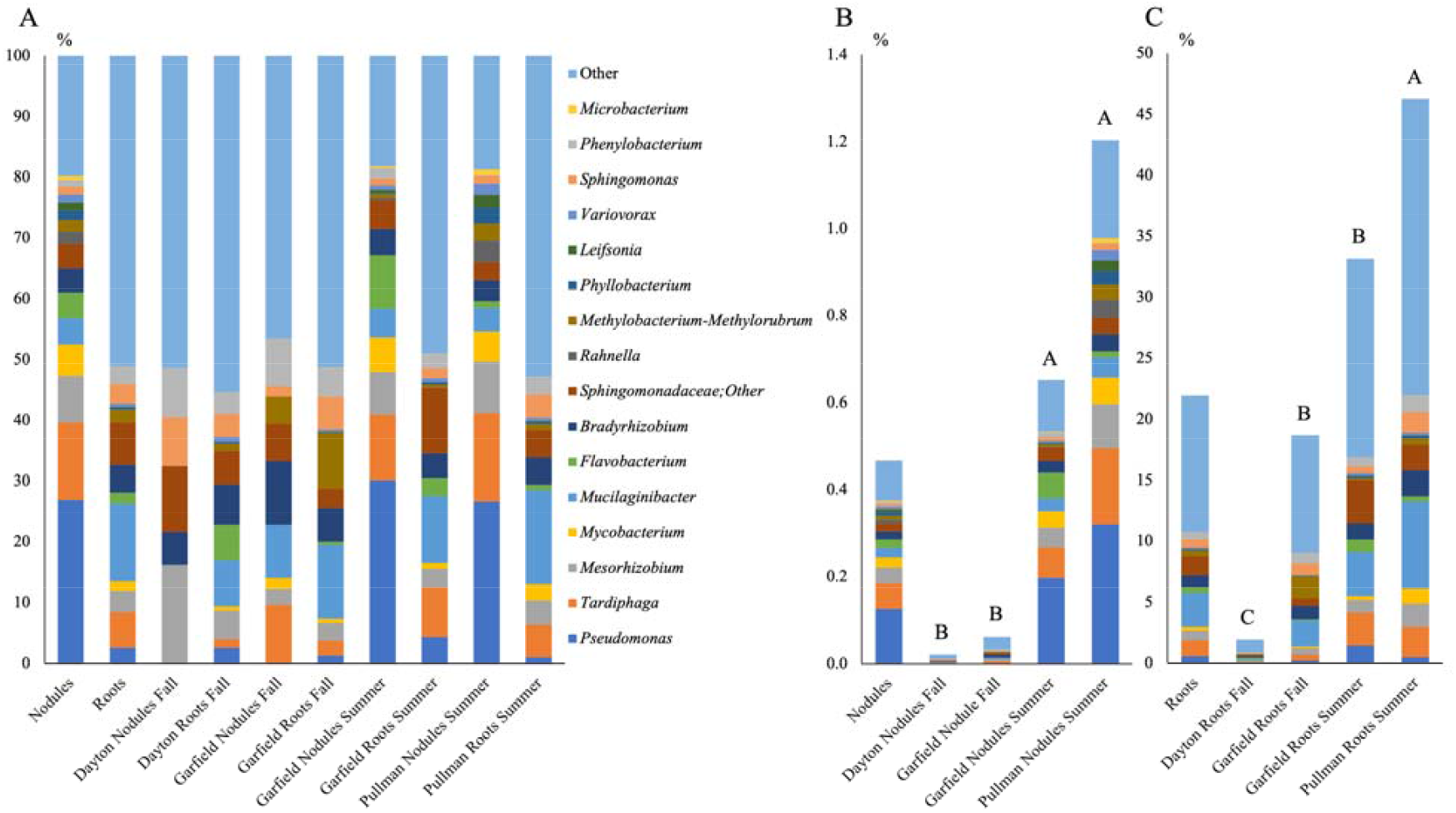
Relative abundances of non-rhizobial taxa detected in the nodule tissue. A – composition of non-rhizobial microbiome; B – proportion of non-rhizobial reads in nodule microbiome, C – proportion of non-rhizobial reads in root microbiome

### 3.4. Site weather conditions and soil chemical properties

Total WP microbiome varied significantly across sampling events (WA fields (location) and collection time). Approximately 13% of the community variation was attributed to these two factors combined. Furthermore, the location influenced bacterial alpha-diversity, which was reflected in significant variation in the Shannon index between sampling sites (Fig. 4), with the community from Dayton exhibiting significantly lower diversity, compared to Pullman and Garfield. The structure of both the nodule and root microbiomes was significantly affected by sampling events; approximately 36% and 27% of the community’s variation was explained by this factor, respectively. The root community collected in Pullman exhibited the highest Shannon diversity (6.3), followed by Garfield Summer (4.3) and Garfield Fall (3.6), with the lowest alpha-diversity found in Dayton root microbiome (Fig. 4). On the other hand, we did not detect substantial variation in the nodule Shannon index between sampling events.

#### 3.4.1. Taxa differentially represented across sampling events

We identified 72 and three bacterial genera in root- and nodule-associated microbiome, respectively, which were differentially represented between sampling events. The relative abundances of 15 most abundant root- and all nodule-associated taxa differentially presented across sampling events are shown on Fig. 7. *Pseudomonas, Tardiphaga*, and *Mesorhizobium* were overrepresented in both root- and nodule-associate microbiomes collected in summer, compared to that in fall. With exception of *Rhizobium* and *Methylobacterium-*Methylorubrum, most other root-associated genera differentially represented across location were more abundant in summer collected samples, compared to the samples collected in fall (Fig. 7).

**Fig. 7.**
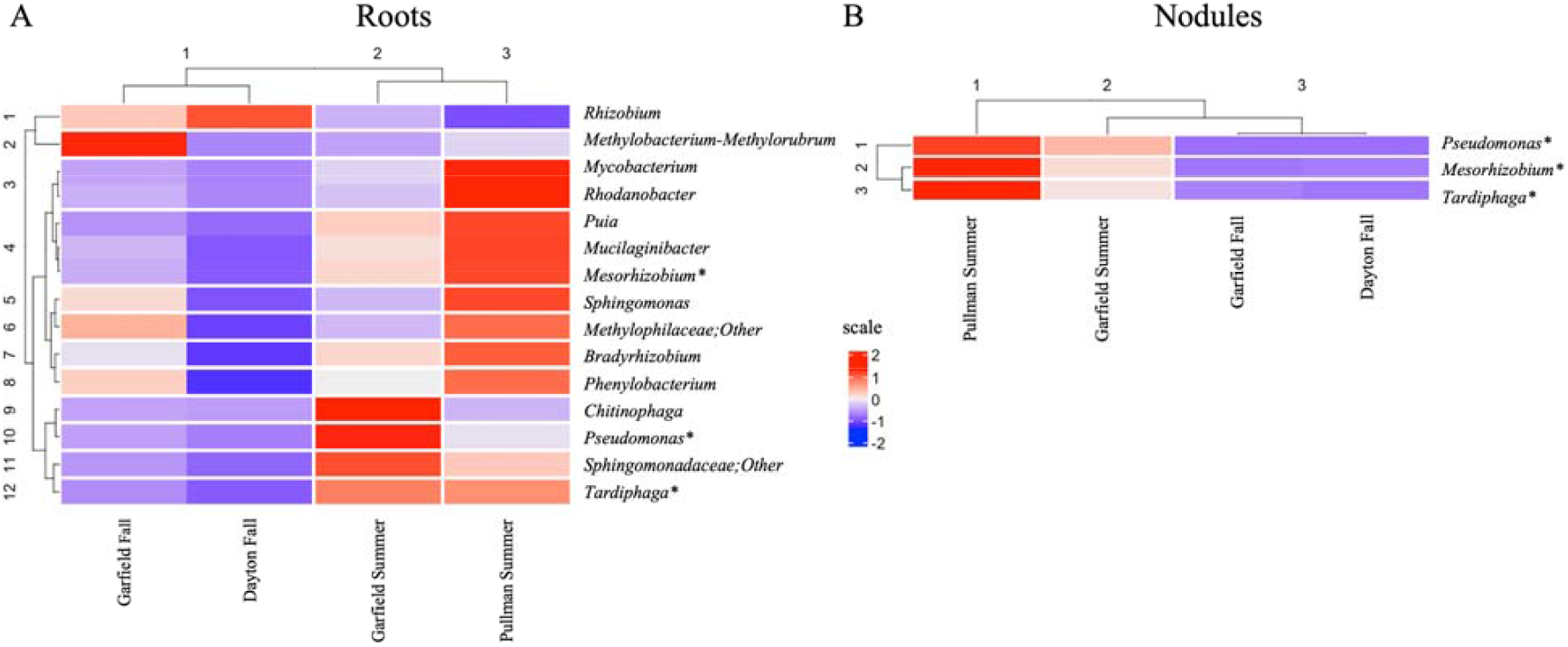
Relative abundances of bacterial genera differentially represented between sampling events. A – 15 most abundant taxa identified in root-associated microbiome; B – bacterial taxa identified in nodule-associated microbiome. Kruskal-Wallis pairwise test (p < 0.05). Corrected p-values were calculated based on Benjamini-Hochberg FDR multiple test correction. * – taxa differentially represented between sampling events in both root- and nodule-associated microbiomes

#### 3.4.2. Rhizobial community

In the combined root- and nodule-associated microbiomes, 32, 108 and 77 ASVs annotated as *Rhizobium* were identified in Dayton, Garfield and Pullman, respectively. In all locations most all rhizobial ASVs found in the nodules were also found in the roots (Fig. 8 A, B, C). However, a significant proportion of rhizobial ASVs were unique to roots: 66%, 61% and 51% ASVs were found only in root samples, from Dayton, Garfield and Pullman, respectively.

**Fig. 8.**
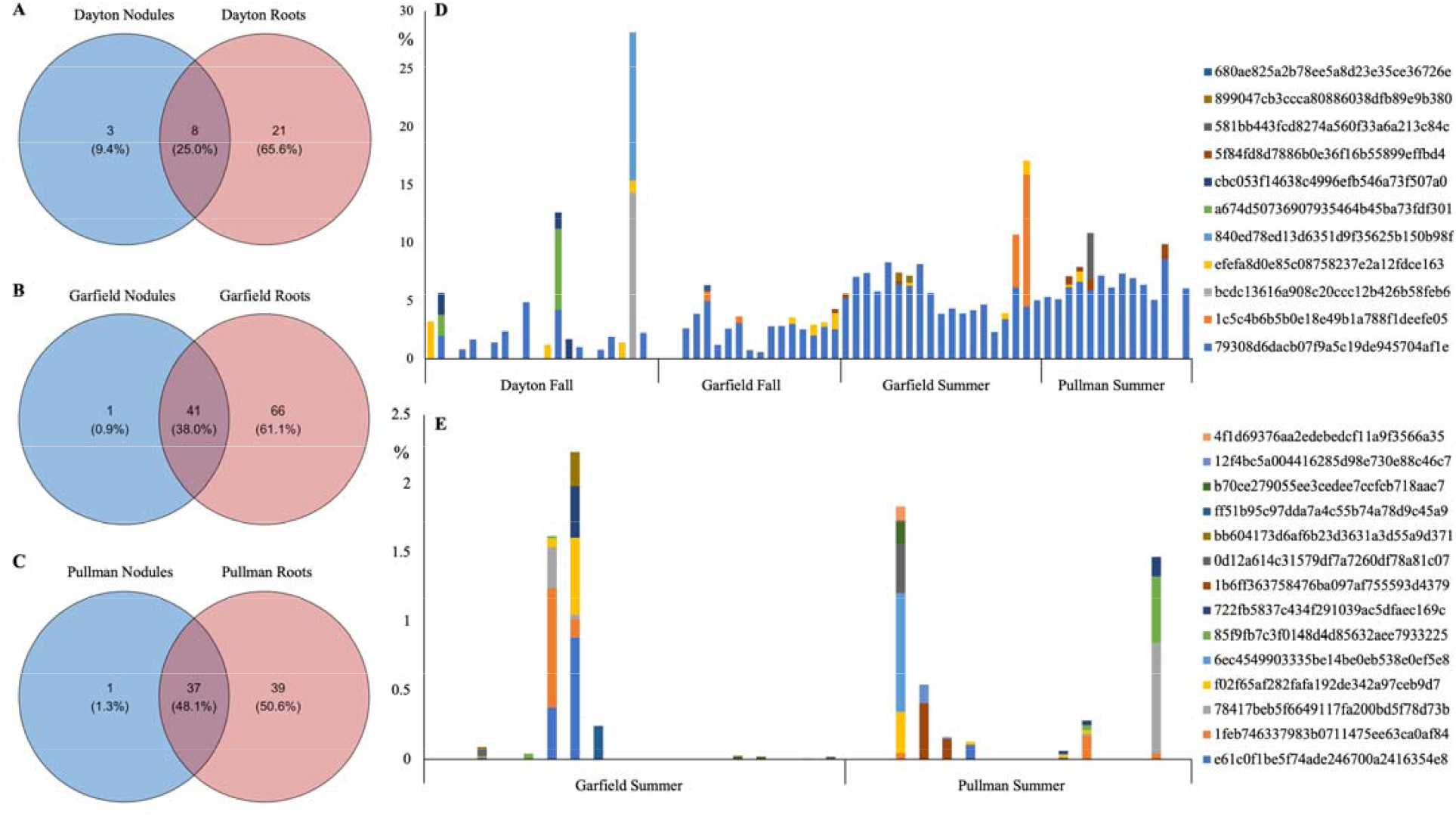
Distribution of *Rhizobium* and *Pseudomonas* ASVs across locations. Venn diagrams showing overlap in rhizobial ASVs between nodule and root tissue in A – Dayton; B – Garfield; and C – Pullman. D – Relative abundances of low abundant *Rhizobium* ASVs across nodule samples, only ASVs represented by at least 0.5% of total nodule reads in at least on samples are listed. E – Relative abundances of *Pseudomonas* ASVs across nodule samples. D and E – The values represent % of the reads in the sample

The ASV1, ASV2 and ASV3, were dominant in the root- and nodule-associated microbiomes in all locations (Fig. 5), and were represented by around 97%, 95% and 92% of total nodule reads, in Dayton, Garfield and Pullman, respectively (Table S3), suggesting potential role of these ASVs in WP nodulation. The ASV1, ASV2 and ASV3 were significantly overrepresented in Garfield’s nodules compared to roots, and ASV2 and ASV3 were overrepresented the nodules from Dayton and Pullman, respectively (Fig. 5). Additionally, ASV2 was significantly overrepresented in Garfield fall-collected nodules, compared to those collected in summer (Fig. S2), suggesting that the rhizobial group associated with this ASV might be more efficient in forming nodules in fall. Both nodules and roots contained a substantial number of rhizobial ASVs unique to the individual locations, around 9%, 36% and 30% of the ASVs identified in total nodule-associated microbiome were unique to Dayton, Garfield and Pullman, respectively (Fig. 2 C). However, these ASVs might be tightly attached to the nodule surface rather than be part of internal nodule microbiome. Additionally, a number of relatively low abundant rhizobial ASVs, with relative abundances >0.5% of reads in at least one sample, were sporadically identified in the nodules (Fig. 8 D; Table S7). Genetic diversity of these ASVs is represented in Fig. S3. Some of these ASVs were unique to the location, while others were found across all locations. For example, ASV bcdc13616a908c20ccc12b426b58feb6 was represented by 14% reads in only one Dayton sample, and ASV 1c5c4b6b5b0e18e49b1a788f1deefe05 was found in four nodule samples collected in Garfield.

#### 3.4.3. Non-rhizobial community

In Garfield (summer and fall collection time) and Pullman, genus Rhizobium was significantly overrepresented in nodule, compared to root tissue (Table S5). However, no difference in Rhizobium between root and nodule tissue was detected in Dayton. While we did not detect any non-rhizobial taxa significantly overrepresented in the nodule-associated microbiomes across individual sampling events, the genus *Pseudomonas* was apparently enriched in the summer-collected nodules compared to roots. For example, summer-collected Garfield’s nodules contained 30% of non-rhizobial reads annotated as *Pseudomonas*, compared to 4% of the reads in the corresponding roots (Fig. 6 A; Table S6). Similarly, in Pullman 27% and 1% of all non-rhizobial reads were annotated as *Pseudomonas* in nodule and root samples, respectively. However, distribution of *Pseudomonas* reads was sporadic – it was detected in less than 30% of nodule samples. For example, no *Pseudomonas* reads were detected in fall-collected Dayton or Garfield nodules (Fig. 6 A; Table S6) and many summer-collected nodule samples did not contain *Pseudomonas* reads (Fig. 8 D; Table S7). Furthermore, there was a remarkable diversity of Pseudomonas ASVs across samples. In total 33 *Pseudomonas* ASVs were identified in the WP nodules, some of which were relatively highly abundant in one or two samples regardless of the location, while some were found in a single sample, and others were detected in relatively low abundance in a few samples (Table S8). Therefore, we could not identify any *Pseudomonas* ASV unique to a specific location or dominant in WP nodules.

We detected a significant difference in the proportion of non-rhizobial ASVs between microbiomes collected in summer and fall. Only 0.02% and 0.06% of the reads were annotated as non-rhizobial ASVs in fall-collected nodules samples in Dayton and Garfield, respectively, while 0.38% and 1.2% of the reads were annotated as non-rhizobial ASVs in summer collected nodule samples in Garfield and Pullman, respectively (Fig. 6 B). Root microbiome collected in Dayton had the least non-rhizobial reads (2% total), while Pullman roots contained the highest number of non-rhizobial reads (46%). There were no significant differences in the number of reads representing non-rhizobial taxa between the fall- and summer-collected Garfield roots.

### 3.5. Soil microbiome

Both location and collection time significantly affected soil microbiome structure. However, location explained 50% of soil community variation, while collection time accounted only for 4% of community variation (Table 1; Fig. 9 A). We also detected significant differences in bacterial alpha-diversity between sampling sites. Dayton’s soils exhibited highest Shannon diversity (9.6 and 9.8 for summer and fall sampling, respectively), followed by Garfield’s (9.3 and 9.1) and Pullman’s (8.8 and 8.4) soils. In Garfield and Pullman, soils collected in summer exhibited significantly higher Shannon diversity compared to those collected in fall, while in Dayton, the collection time had no significant effect on the Shannon diversity (Fig. 9 B). We identified 147 bacterial genera differentially represented between locations (Table S9). However, this list did not include either *Rhizobium* or *Pseudomonas*. On the other hand, few changes in bacterial community composition were detected between sampling times. Only two relatively low abundance taxa, uncultured Microscillaceae and Planctomycetota vadinHA49, were overrepresented in summer collected soils, compared to ones collected in fall (Table S10). *R. leguminosarum* ASV1, ASV2, ASV1, comprising most of rhizobial ASVs in the nodule microbiomes were sporadically detected in the soil (Table S11). ASV1, ASV2, and ASV3 were detected in all soil samples, albeit at low abundance (26, 30, and 55 reads, respectively).

**Fig. 9.**
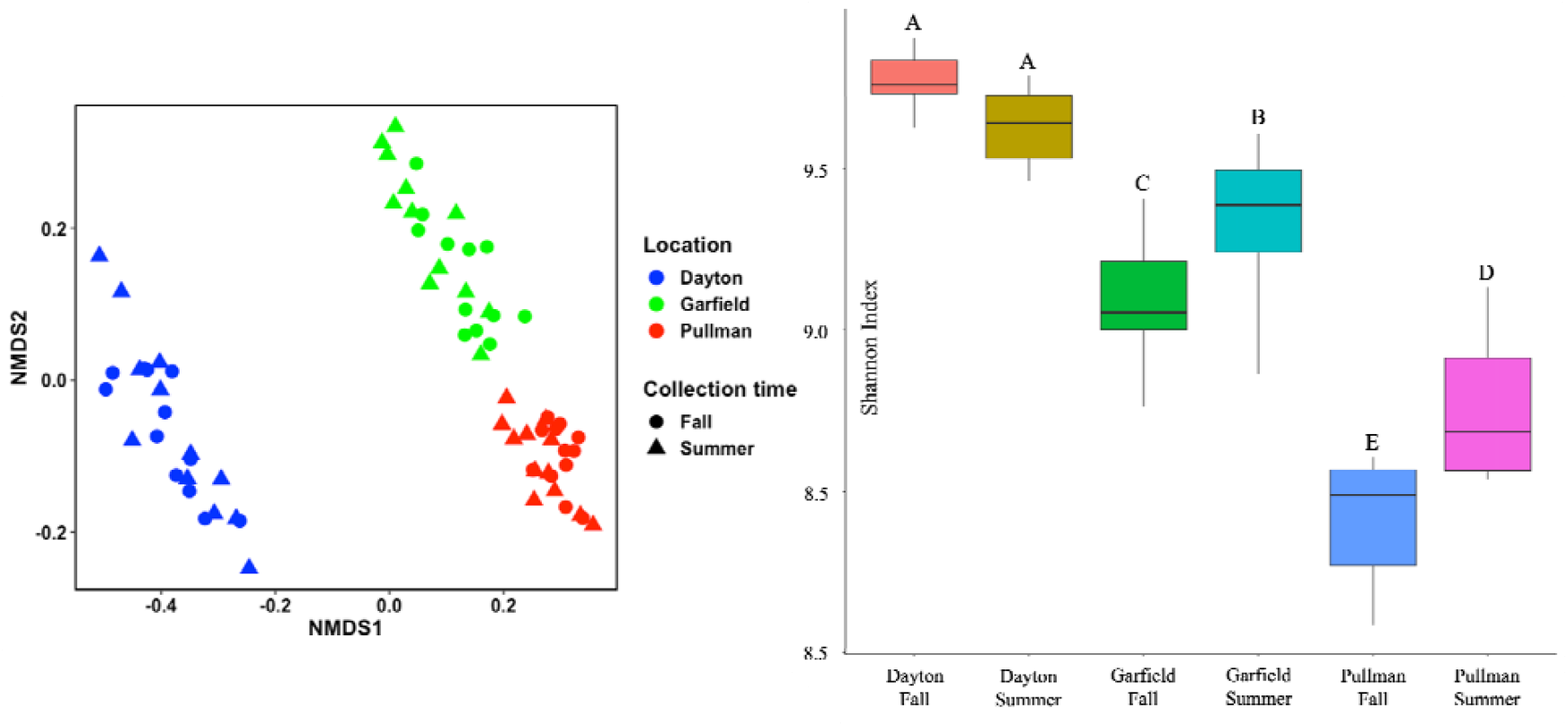
Soil microbiome. A - Nonmetric multidimensional scaling (NMDS, based on Bray-Curtis dissimilarity distances) of bacterial communities at ASV level. B – Estimated Shannon diversity of soil bacterial community. For each variable, data followed by different letters are significantly different according to Kruskal-Wallis pairwise test (p < 0.05). Corrected p-values were calculated based on Benjamini-Hochberg FDR multiple test correction

## 4. Discussion

A standard approach in studying plant-microbe interaction implies consideration of the plant as a host for the associated microbiome. However, legume nodules represent a unique niche where host-plant and host-associated rhizobia must be considered as partners hosting non-rhizobial microorganisms. The nodule environment is strikingly different from nodule-free root tissue in both its metabolic and genetic structure. A nodule’s function requires metabolic cooperation between legume and rhizobia to provide nitrogen fixing capability. This involves highly attenuated nitrogen and carbon transformation and utilization, such as amino acid and dicarboxylic acid metabolism, coupled with N-fixation and tight control of oxygen availability (Basile & Lepek, 2021). Single nodules can contain up to 10^9^ rhizobial cells, contributing dramatically to the nodule gene pool (Downie, 2014). In agreement with this, our data indicate that the nodule microbiome differs significantly from that of the root. This was evidenced by the differences between root and nodule overall community structures and alpha-diversity. The decrease in nodule Shannon diversity could be explained by overpopulation of the nodules with a limited number of nodule-induced rhizobia (Gage, 2002). Indeed, *Rhizobium* was represented by 99.5% of total reads in the nodule microbiome, in contrast to 78% of reads in the root microbiome. More specifically, *R. leguminosarum* ASV1, ASV2 and ASV3 were represented by 95% and 63% of total nodule and root reads, respectively.

As WP is planted in fall, there is a potential for the legume to develop early nodules by late November – early December. Therefore, we assessed WP nodulation in December 2021 in three locations across WA and detected nodule formation in two locations, Dayton and Garfield, but not in Pullman. In controlled laboratory and green-house conditions, germinated pea seedlings usually develop nodules within two weeks. The similar trend was observed in our plant test performed in laboratory conditions. However, in field conditions, especially in cold fall weather, it takes longer for seeds to germinate and may take longer for rhizobia to colonize plant roots and trigger nodule development. The plants in Pullman were very small with mostly primary roots and no apparent nodules. The delay in plant development might be a result of combined effect of local climate, soil conditions and the time of WP planting. We did not detect any substantial differences in overall weather conditions, such as soil and air temperatures, between Garfield and Pullman in November. On the other hand, in November, Pullman received substantially higher level of precipitation, compared to Garfield and Dayton, resulting in waterlogged soil. However, because of the limited data points, we cannot make any strong conclusion regarding factors affecting WP early nodulation. Most probably the factors affecting plant development might play a significant role in the early nodulation event. Nevertheless, the question of the ability of WP to form nodules in fall and the role of the early nodulation in plant fitness and production is important and should be addressed.

The effect of plant genotype on root- and nodule-associated microbiome was one of the questions we addressed in this study. Our data indicate that cultivar did not affect the WP root and nodules microbiome profile, since we did not identify any specific taxa differentially represented between cultivars, including rhizobium species. However, plant genotype was a significant factor affecting the overall structure of nodule microbiome. Similarly, a previous field study of the symbiotic preference of white clover for native rhizobial populations indicated that while genetically distant clover crosses did not exclusively select for specific *R. leguminosarum* alleles, plant genotype was a strong factor affecting nodule rhizobium composition (Fields *et al*., 2023).

The WP microbiome was significantly affected by sampling event. Total microbiome associated with the plants grown in Dayton exhibited the lowest Shannon index, compared to that from Garfield and Pullman, which might be partially explained by the fact that all Dayton samples were collected in fall and contained plant tissue at a relatively early developmental stage. However, we did not detect a significant decrease of Shannon index in fall-collected microbiomes in Garfield, compared to that in summer-collected microbiomes. A small variation in nodule bacterial diversity was detected between locations, which could be also explained by rhizobial dominance. In contrast to nodule-associated microbiomes, diversity of root microbiomes differed significantly between the locations, with Dayton samples having the lowest and Pullman samples having highest Shannon diversity. Similarly, a significant role of local environmental conditions in nodule rhizobium composition was reported in a white clover field study (Fields *et al*., 2023). Next, we tested whether the differences in root-associated microbiome alpha-diversity could be explained by the variations in soil microbial communities. Interestingly, soil bacterial diversity exhibited an opposite trend to that of plant-associated microbiomes. This suggested that other factors might play a role in diversification of WP root bacteria. On the other hand, soil chemical characteristics and weather conditions could affect the soil microbiome. For example, previously we showed that soil pH was one of the most important variables affecting bacterial community structure (Yurgel *et al*., 2018). On average the soil and air temperatures and soil pH were higher in Dayton than in Garfield and Pullman, and could explain the higher bacterial Shannon diversity in Dayton soil.

Root-associated microbiomes provided a foundation for the nodule bacterial communities. The majority of ASVs found in the nodules were also found in the plant roots. Nevertheless, nearly half of rhizobial ASVs were found only in root tissue. Additionally, the nodule microbiome was dominated by three *R. leguminosarum* ASVs, that represented 80% to 100% of total nodule reads across all samples and locations. These ASVs were represented by the majority reads in the root tissue. We also detected a few rhizobial ASVs specific to the individual locations. Some of these rhizobia were identified in relatively high abundances in one or two nodule samples, suggesting that they might be part of more intimate symbiotic interaction with the host plant, rather than being tightly attached to the nodule surface. These data indicated that (i) there was a large population of soil derived rhizobia that were strongly attracted by WP roots; (ii) a smaller proportion of these rhizobia was also a part of the nodule-associated microbiome; (iii) there were a few highly abundant rhizobial groups overrepresented in the nodule-associated microbiome and common to all tested locations, suggesting their potential involvement in WP nodulation; and (iv) there was a limited number of potential symbionts unique to the individual locations. It is important to emphasize that each nodule sample was derived from several nodules from either a single plant or from several plants combined, and therefore it does not represent the community of a single nodule. Additionally, 16S rRNA amplicon sequencing might not provide a strong taxonomic resolution at strain level (Johnson *et al*., 2019), especially considering genetic diversity of *R. leguminosarum* (Kumar *et al*., 2015).

There are numerous studies reporting identification of non-rhizobial nodule residents (Ibáñez *et al*., 2009, Ferchichi *et al*., 2019, Crosbie *et al*., 2022, Martínez-Hidalgo & Hirsch, 2022, Hossain *et al*., 2023). However, there are few circumstances that indicate that these non-rhizobial microorganisms were not just a part of the “normal” root microbiome but were directly selected/enriched by legume-rhizobia host for nodule occupancy. Here we attempted to address this question by focusing on non-rhizobial WP nodule microbiome. The major obstacle for the identification of non-rhizobial microorganisms enriched in legume nodules using next-generation sequencing is an overpopulation of rhizobia in the nodule tissue. Since the abundances of microorganisms estimated by amplicon sequencing are relative and limited by the sequencing depth of the samples, the overpopulation of a community by a few ASVs dramatically decreases relative abundances of other taxa. For example, in the root microbiome, 22% of the total reads represented non-rhizobial ASVs, compared to 0.5% of that in nodule microbiomes. Therefore, a direct comparison of root and nodule microbiome, with the goal to identify differentially represented taxa, would ultimately point to a significant decrease in relative abundances of most non-rhizobial microorganisms in the nodules compared to roots. This was what we detected in our analysis. We attempted to overcome this obstacle by considering both plant and rhizobial reads as “host” sequences and filtered them out. However, this approach produced a very limited subset of samples containing just 12% and 87% of original number of nodule and root samples, respectively. Additionally, tissue of fall-collected nodules contained significantly less non-rhizobial bacteria, compared to summer collected nodules, and only a few summer-collected nodules retained >100 non-rhizobial reads. This indicated that colonization of nodule tissue by non-rhizobial bacterial occurred in later stages of nodule development, following rhizobial plant cell colonization and nodule tissue expansion, probably caused by prolonged exposure to the diverse soil microbial community and potentially nodule senescing. Because of these limitations we did not identify any non-rhizobial taxa significantly overrepresented in nodule compared to root tissue. However, a future study focusing on mature N-fixing nodules with increased depth of amplicon sequencing might provide better resolution of nodule specific non-rhizobial nodule residents.

It is well-known that *Pseudomonas* can promote plant growth under environmental stresses (Rajkumar *et al*., 2017). Several previously published studies reported inter- and intra-cellular presence of *Pseudomonas* in the legume nodules (Hakim *et al*., 2020, Hansen *et al*., 2020, Mayhood & Mirza, 2021, Pastor-Bueis *et al*., 2021, Crosbie *et al*., 2022, Hossain *et al*., 2023). Recently it was shown that *Pseudomonas* can selectively colonize healthy plant nodules and promote establishment of effective symbiosis in *Lotus*□*japonicus* (Crosbie *et al*., 2022), while co-inoculation of *Pseudomonas* with *Mesorhizobium* led to an increase in nodule number in chickpea (Malik & Sindhu, 2011). On the other hand, *P. syringae* and other species belonging to the *P. syringae* group can cause severe damage to many plant species (Tribelli & López, 2022, Yang *et al*., 2023). *Pseudomonas* was the most prevalent non-rhizobial taxon in the nodule-associated microbiomes and was apparently enriched in the summer-collected nodules compared to roots. The presence of this taxon in the nodule-associated microbiome might be partially attributed to their colonization of senescent zones IV.

In summary, this study showed that WP cultivars are capable of forming nodules in PNS soils in late fall. However, the extent of the nodulation was not uniform across the locations, suggesting a strong effect of environmental factors on this process. The role of WP early nodulation in crop production and resistance to environmental stresses is an important question that should be further investigated. Our data also indicated that a diverse population of native rhizobia can colonize WP roots cultivated in WA soils. However, a substantially smaller subset of these bacteria is able to colonize WP nodules. Interestingly, three ASVs were dominant nodule-associated rhizobia in all tested locations regardless of the significant variation in soil microbiome diversity and structure between locations. These ASVs had relatively low abundance in the soils, indicating their strong attraction to host-plant roots and nodules. Our data also indicated an enrichment of WP nodule-associated community collected in summer with a few non-rhizobial bacterial taxa, compared to that collected in fall. Finally, while the complementation of legume nodule microbiome study with root-associated microbiome analysis might be a useful tool to identify non-rhizobial nodule residents, future study focusing on mature N-fixing nodules with increased depth of amplicon sequencing might provide a better resolution of nodule specific non-rhizobial nodule residents.

## Abbreviations

WP: winter pea
WA: the US state of Washington
N: nitrogen
PNW: Pacific Northwest

## Author Contributions

Conceptualization, S.N.Y and R.M.; methodology, S.N.Y and R.M.; formal analysis, S.N.Y.; writing—original draft preparation, S.N.Y.; writing—review and editing, S.N.Y. and R.M. All authors have read and agreed to the published version of the manuscript.

## Data Availability Statement

The datasets generated for this study can be found in the NCBI se-quence read archive under the accession numbers PRJNA1073214 and PRJNA1073218.

## Acknowledgments

Jarrod Pfaff and Jeff Haines are thanked for planting and maintaining the field plots. Weather data provided courtesy of Washington State University AgWeatherNet. Data are copyright of Washington State University.

---

**Figure S1.**
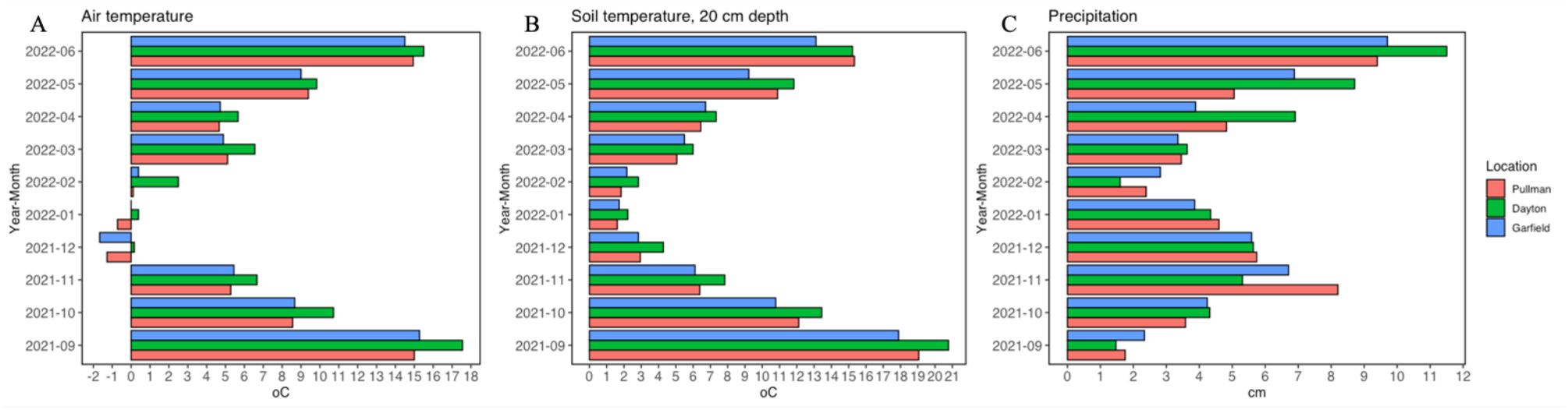
Weather conditions In the sampling site. A – Average air temperature, ° C; B – Average soil temperature at 20 cm depth. ° C; C – Total precipitation per month, cm.

**Figure S2.**
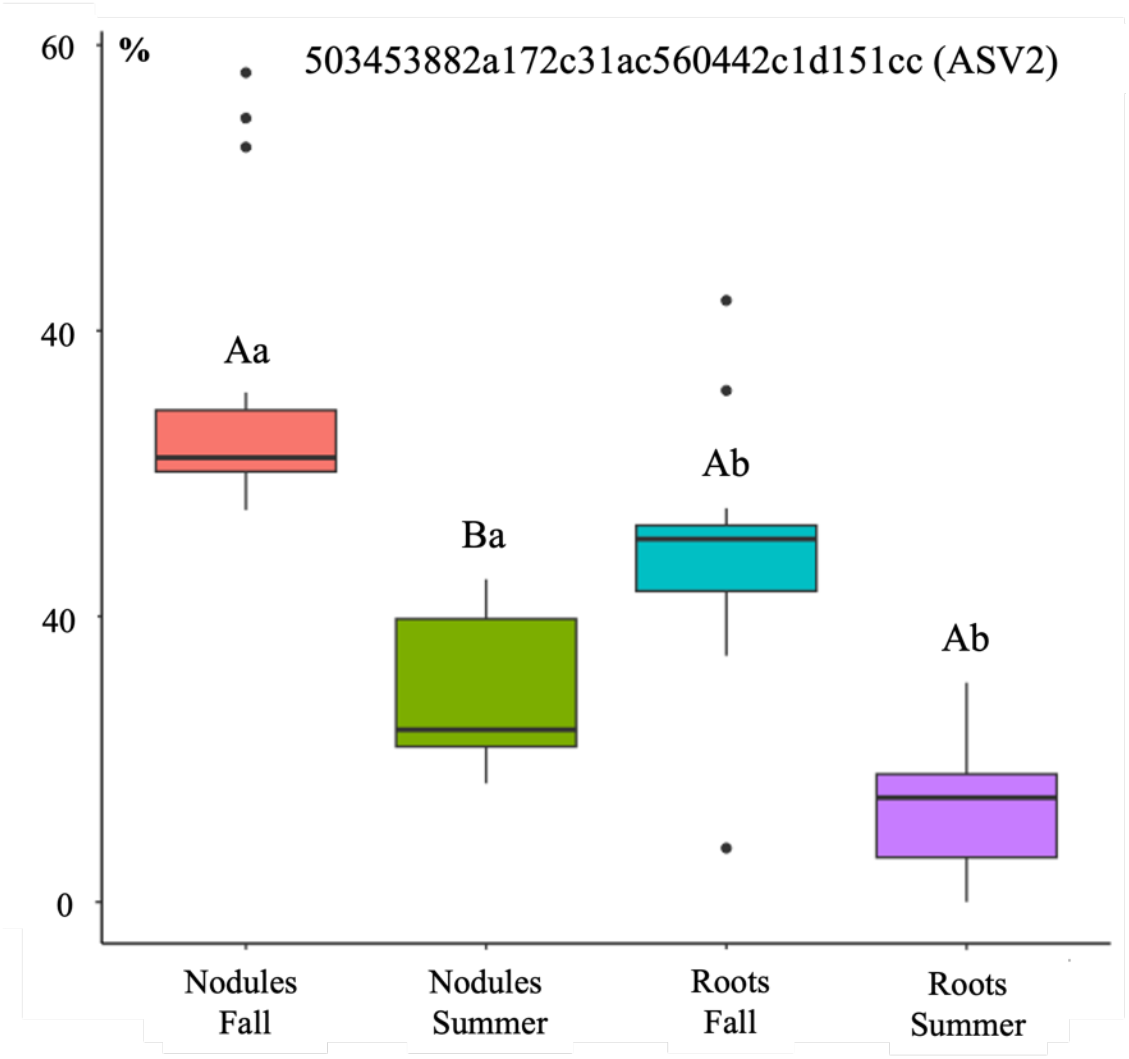
Relative abundance of ASV2 in Garfield microbiomes. For each variable, data followed by different letters are significantly different according to Kruskal-Wallis pairwise test (p < 0.05). Corrected p-values were calculated based on Benjamini-Hochberg FDR multiple test correction. Capital letters indicate differences between collection time within same tissue. Lowercase letters indicate differences between tissue type within same collection time.

**Figure S3.**
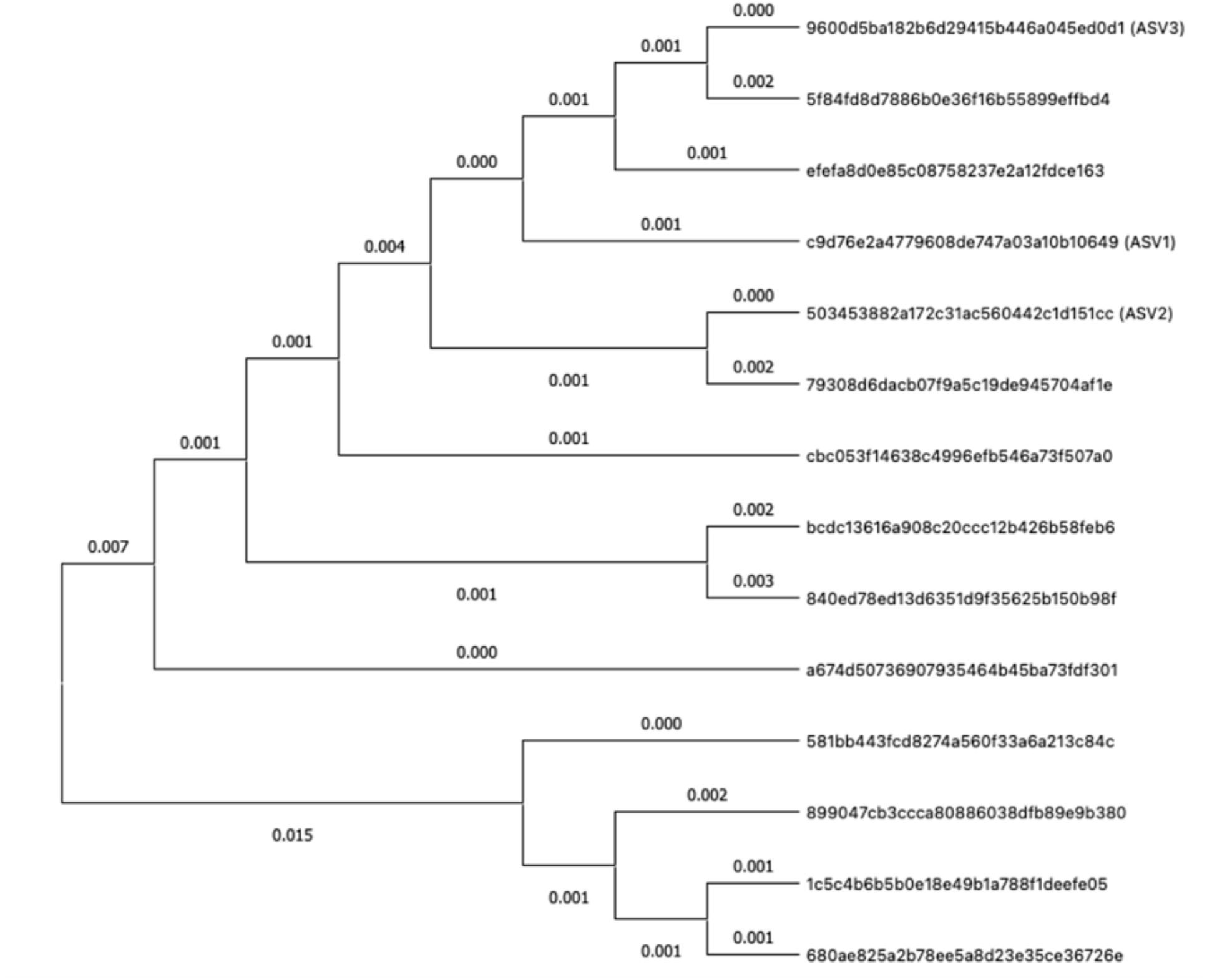
Genetic diversity of *R. leguminosarum* ASVs. The evolutionary history was inferred using the Neighbor-Joining method [1, 2]. The optimal tree is shown (next to the branches). The evolutionary distances were computed using the Maximum Composite Likelihood method [3] and are in the units of the number of base substitutions per site. This analysis involved 14 nucleotide sequences. All ambiguous positions were removed to reach sequence pair (pairwise deletion option). There was a total of 418 positions in the final dataset. Evolutionary analyses were conducted in MEGA11 [4,5]. [1] Saitou N. and Nel M. (1987). The neighbor-joining method: A new method tor reconstructing phylogenetic trees. Molecular Biology and Evolution 4:406-425, [2] Felsenstein J. (1985). Confidence limits on phylogenies: An approach using the bootstrap. Evolution 39:783-791, [3] Tamura K., Nei M., and Kumar S. (2004). Prospects for inferring very large phylogenies by using the neighbor-joining method. Proceedings of the National Academy of Sciences (USA) 101:11030-11035, [4] Tamura K., Stccher G.. and Kumar S. (2021). MEGA 11: Molecular Evolutionary Genetics Analysis Version 11. Molecular Biology and Evolution https://doi.org/10.1093/molbev/msabl20, [5] Stecher G., Tamura K., and Kumar S. (2020). Molecular Evolutionary Genetics Analvsis (MEGA) for macOS. Molecular Biology and Evolution 37:1237-1239

